# Protein structure featurization via standard image classification neural networks

**DOI:** 10.1101/841783

**Authors:** Tobias Sikosek

## Abstract

Many applications in the biomedical domain involve the detailed molecular and functional characterization of macro-molecules such as proteins. Where possible, this involves the knowledge of detailed 3D coordinates of every atom within a protein. At the same time, machine learning has become the basis of much innovation within this domain in recent years. There are, however, a few challenges in applying machine learning to 3D protein structures, such as variability in size and high dimensionality of the data. It would therefore be beneficial to be able to map every protein structure to a smaller fixed-dimensional representation that is directly learned from the structure without manual curation. In addition, it would be valuable for biomedical researchers if such approaches would require little method development and instead draw from cutting-edge research such as image classification via deep neural networks. Here, such an approach is outlined that first re-formats protein structures as 2D color images and then applies off-the-shelf neural networks for image classification. It is shown that such neural networks can be trained to effectively encode the CATH protein classification database and that feature vectors extracted from such networks, once trained, can be transferred to a completely new task that is likely to benefit from molecular protein information, namely that of small molecule binding.

## INTRODUCTION

Understanding protein structure and function is of interest in a wide range of Biological applications. One application that is of most direct relevance to human health is the implication of various proteins in diseases and the role that these proteins may play in finding a therapy.

While many experimental and computational techniques exist to study proteins at detail there is typically a trade-off with the throughput of that same method. High-throughput methods typically come at the cost of molecular detail. This is certainly true for all methods that are used to understand the molecular structures of proteins, which are complex polymer chains of typically 20 amino acid building blocks that fold up on themselves to produce often highly specific three-dimensional arrangements or folds. These folds are then used to endow the protein with a specific biochemical function as part of healthy Biology, but also in the context of disease. Computationally, proteins have been studied at the amino acid sequence level through comparative Bioinformatics (high throughput) or at the biophysical level through Molecular Dynamics simulations of their 3D structures (low throughput). A middle ground between these two approaches is reached in Structural Bioinformatics where the known protein structures are compared based on a combination of sequence and structure-based descriptors so that highly similar proteins receive a high score. These methods calculate a similarity or difference score of pairs of proteins. This score requires that the proteins are assigned to a set of descriptors that allow for a comparison in some proxy feature space that is used to capture actual protein differences. All such descriptors are typically manually selected by the method developer and thus are limited to preconceived notions of what information is important for representing and comparing proteins.

Machine learning builds models by processing large amounts of training samples and optimizing a set of internal parameters. These parameters are fine-tuned so that the model can make predictions on a provided ground truth data label, based on the input features or descriptors of the modeled entities.

Proteins are challenging for machine learning because of their complex 3D structures of variable sizes and the complex relationship between features and prediction targets. However, promising new approaches can now be developed due to the advances in Deep Learning where multi-layer artificial neural networks can be used to extract such complex relationships.

The amino acid sequence of the protein has been used for machine learning by exploiting the similarities to Natural Language Processing. One approach, called ProtVec, has applied word embeddings to amino acid triplets as the “words” and the protein sequence as the “sentence”[1]. However, the sequence of a protein does not specify the biologically active 3D structure – a well-known problem in biology [2].

The most common approach of applying Deep Learning to protein structures is through the discretization of 3D space into bins fed into Deep Convolutional Neural Networks (CNNs) as 3D arrays [3]–[5]. Typically, this requires the definition of a 3D box with a given size and granularity (size of bins) centered on a region of interest such as a known ligand binding site. While this representation of protein structures is very intuitive and interpretable, it comes with some down-sides such as a high degree of sparseness and dimensionality as well as computational costs resulting from 3D input data. Additionally, the development of methods for 2D images generally proceeds more rapidly than that for 3D approaches, so that the adoption of existing deep learning techniques is easier in 2D.

A different route has been taken here, where the protein structure is not interpreted as a set of absolute 3D atomic positions, but rather as a set of pairwise distances. This effectively allows to treat the protein structures as flat, image-like 2D arrays. Such distance matrices have two identical axes corresponding to an ordered list of atoms starting from amino acids at the N-terminal end of the protein backbone and ending at the C-terminal end. Atoms of the same amino acid are typically listed in a standard order, for example in the PDB format of the Protein Data Bank, allowing the formation of predictable patterns. Recently, a few studies have been published that also follow this general approach, but only consider a single atom per amino acid and use their methods for protein search [6] and protein design [7] applications. For the application of drug discovery, all-atom molecular detail will be needed in the future, although the prototype presented here does not yet achieve that level of detail.

Here, we focus on the case where the category that we want to predict is the ability of a protein to bind a small molecule such as a drug, or not. Such drug molecules are designed to bind to certain proteins, typically to alter the behavior of that protein in a disease context and with the aim of a therapeutic effect.

In the first part of this work, the focus lies on training a neural network on protein structure classification in general, and without a specific application in mind. This is to maximize generalizability of the protein features and avoid hidden biases of small molecule activity datasets. The results clearly show that protein structure classification can be solved by Deep Learning without much difficulty.

In the second part, protein feature vectors (fingerprints) generated based on the trained model from the first part are applied to an activity data set where the strength of an interaction between a small molecule and a protein was measured experimentally.

## METHODS

### Dataset preparation

#### CATH

The CATH [8] non-redundant data set was used as a training set for protein fingerprint generation. These include 21090 3D domain structures from across the entire CATH classification system and constitute a small but less biased subset of the full 308999 domain structures. Data labels for supervised learning were taken from the ‘Class’, ‘Architecture’, ‘Topology’, and ‘Homologous superfamily’ levels with 4, 40, 1364, and 2714 sub-categories, respectively. Note that these numbers are partially slightly lower than the ones published on CATH, due to loss of some protein structures during processing as outlined further below.

#### ChEMBL

ChEMBL [9] version 23 was used as a starting point to select activity values of small-molecule ligands and their single protein targets for which a pChEMBL value was available. pChEMBL is defined as: -Log(molar IC50, XC50, EC50, AC50, Ki, Kd or Potency), depending on the assay. Based on the pChEMBL value, three activity categories were defined that approximately balanced the number of low and high activity observations, while keeping the intermediate activity bin relatively small. This was obtained by choosing pChEMBL values of 5.2 and 6.0 as the lower and upper bounds of that intermediate bin, respectively. Furthermore, the data set was restricted to cases where each compound and each target was associated with at least one high-activity and one low-activity observation, thus arriving at a final dataset of 35076 compound and 871 targets, with a total of 218635 activity measurements. ChEMBL single protein targets are associated with a Uniprot identifier corresponding to the underlying gene/protein of that target. For each Uniprot identifier, multiple 3D structures (mostly from X-ray crystallography and obtained from the Protein Data Bank, PDB) can be found that may correspond to different structural subunits of that protein. To automatically match each ChEMBL target with a protein structure, the target protein sequence reported by ChEMBL is compared to the sequence of all PDB structures (chains) linked to the same Uniprot ID. The PDB structure with the highest pairwise alignment score, calculated with BioPython (https://biopython.org/), was used as that target’s structure – as long as a minimum score of 50 was obtained (parameter-free Bio.pairwise2.globalxx function). Otherwise, the target was discarded. Among alternative structures for the same target with identical alignment score, the one with the lower resolution value was picked. Any remaining ties were resolved by picking the structure with smaller ligand size, to reduce the conformational bias towards the crystallized ligand.

### Protein structure to image conversion

All protein structures used were converted to a format that could be used directly as input for state-of-the-art Deep Convolutional Neural Networks. The three channels typically reserved for the Red, Green, and Blue (RGB) color contributions of images, were populated with three different 2D representations of a single protein structure: 1) the pairwise atomic distances of heavy atoms in the protein structure (D); 2) a pairwise non-bonded energy term like the one used in the Amber molecular mechanics force field [10] (NB); 3) the atomic cross-correlation matrix from an anisotropic network model (ANM)[11], [12]. ANMs are used to calculate the normal modes of protein structures, i.e. their intrinsic motions and dynamics. The equation for the NB channel was of the standard Amber [13] form where E_ij_ is the potential energy contribution of interactions between atoms i and j, with partial charges q_i_ and q_j_, as well as actual distance r_ij_ and equilibrium distance r^0^_ij_:

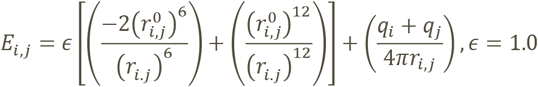

Distances D were cut off at an upper limit of 51.2 Å whereas no lower limit was applied. ANM correlation matrices range between −1 and 1. The ProDy [14] python library (http://prody.csb.pitt.edu/) was used to calculate D and ANM. NB was calculated based on the partial charges assigned by the PDB2PQR tool (http://www.poissonboltzmann.org/pdb2pqr/) using the ‘Amber’ force field option and cut off outside a range of (−0.1, 0.1).

After these processing steps, 20798 domain structures remained in the final CATH data set, where some PDB structures caused errors and were thus discarded.

Finally, each RGB image was down-scaled to 224×224 pixels for DenseNet121, ResNet50, VGG16, U-Net, and 299×299 pixels for Inception v3.

### Deep learning on protein structures

We used the VGG16 [15], Inception V3 [16], ResNet50 [17], and DenseNet121 [18] architectures provided by Keras (https://keras.io) (GPU optimized version 2.1.5, using Tensorflow 1.7 as backend) and with pre-trained weights from ImageNet [19], but without their original softmax output layers. Instead, each network was equipped with one new dense (fully connected) layer using batch normalization, ReLU activation, and Dropout (with a fraction of 0.25 neurons randomly turned off at each training iteration). This dense layer either had 128 or 512 neurons as described in Results, depending on which length feature vector was desired. This layer was then connected to four separate softmax classification outputs for each of the four CATH levels (see above). Networks were trained with a total batch size of 128 split across four NVIDIA GeForce GTX 1080 Ti GPUs, using the Adam optimizer with a learning rate of 0.001 and categorical cross-entropy as loss function for all four outputs (each contributing equally to the final loss). Learning rates were reduced on plateaus of 5 epochs of stagnating validation loss by a factor of 0.2 towards a minimum learning rate of 0.0001 and early stopping was applied whenever the change in validation loss was lower than 0.001 for 20 epochs. 40% of the data were used for validation during training and training data is shuffled before each epoch. In addition to these supervised classification models, the U-Net [20] architecture was modified to be trained on reconstruction loss, thus effectively serving as an autoencoder with skip connections – as opposed to its original application of providing binary segmentation masks for biomedical images. The outputs of the bottleneck layer of 1024 convolutional filters was passed through global average 2D pooling, thus resulting in a protein feature vector of length 1024. Note, that hyperparameters were not systematically optimized and thus only represent one working solution.

### ChEMBL activity prediction

Small molecule activities were predicted as a binary classification task where each pair of compound and target was assigned to a low activity (‘low’) or high activity (‘high’) class. The classifier was a Random Forest with an ensemble size of 100 and without restricting the depth of trees.

Various strategies for splitting the data set into training and validation data and cross-validation were investigated. Random splits randomly assigned 40% of the data to the validation set, whereas stratified approaches balanced data across the two prediction classes (high vs low). Five-fold cross-validation was used throughout. Group-based cross validation schemes trained on n-1 sub-groups and validated on the remaining group that was not used during training. Groups were defined by k=5 clusters obtained through K-means clustering of the protein fingerprints.

Protein fingerprints were obtained as described above for the 871 protein targets in the final ChEMBL data set. Fingerprints were always obtained as the last ReLU activations of the second fully connected layer, just before the softmax output layers. These activations were obtained by feeding the RGB images of ChEMBL-target protein structures into the convolutional neural network trained on CATH data.

## RESULTS

### Protein structures across folds can be encoded as fixed-dimensional fingerprints

Protein structures can be represented as sets of 3D atomic coordinates in Euclidean space that are connected by covalent bonds to form the universal backbone and side chain organization which is often described as a sequence of amino acid letters. Furthermore, atoms can form close physical connections via non-covalent bonds (H-bond, van der Waals, and electrostatic interactions) even when non-adjacent along the sequence and it is these spatial structural arrangements that give proteins their specific functional properties.

To train a neural network to consistently encode any protein structure into a fixed-length latent space representation, it will need to perform either unsupervised or supervised learning. The former will likely take the form of an autoencoder or related algorithms and condenses the information contained in its inputs automatically without any further input and relying on the reconstruction error for learning. Supervised learning on the other hand requires expert labeled data and performs either classification or regression where the error between predictions and ground truth labels is minimized. The availability of labeled data can be a limitation in many machine learning applications, however, in the case of protein structures, abundant labels are already available in databases such as CATH.

The CATH database curates the structural relationships among all known protein structures and does so by breaking proteins down into subunits called domain which roughly correspond to evolutionarily reused building blocks of larger proteins that may have a high likelihood of folding independently. While the entire dataset consists of around 300,000 domain structures, there is also a much smaller non-redundant dataset of around 21,000 structures that have been selected to cover all varieties of structure without being biased towards those that are highly abundant in PDB. It is therefore a suitable dataset for teaching a deep neural network about what proteins look like, including those that are in principle interesting as drug targets due to their readiness to form stable structures during crystallization and subsequent X-ray diffraction analysis – the source of most structures in PDB. These domains are then categorized at various levels of structural detail, starting at the four ‘Classes’ of proteins with mostly alpha-helical secondary structure content, mostly beta-sheet content, or proteins with both alpha-helix and beta-sheet. The fourth Class is reserved for proteins without clear assignment such as intrinsically disordered proteins. Below this top level follows ‘Architecture’, where the four Classes are further subdivided into a total of 40 categories, followed by the ‘Topology’ level (1373 categories), and ‘Homologous Superfamily’ level (2737 categories).

In order to make use of the state-of-the-art power of deep learning models, each CATH domain was converted to a 2D color image by plotting the inter-atomic distances between heavy atoms that typically make up proteins (i.e. Carbon, Oxygen, Nitrogen, Sulphur – but omitting Hydrogen) and augmenting those distances with additional information stemming from pairwise interactions of partial electric charges between atoms as is commonly used in Molecular Mechanics force fields such as Amber, as well as correlated atomic motions predicted by a normal mode calculation (see Methods and Figure 1).

**Figure 1:**
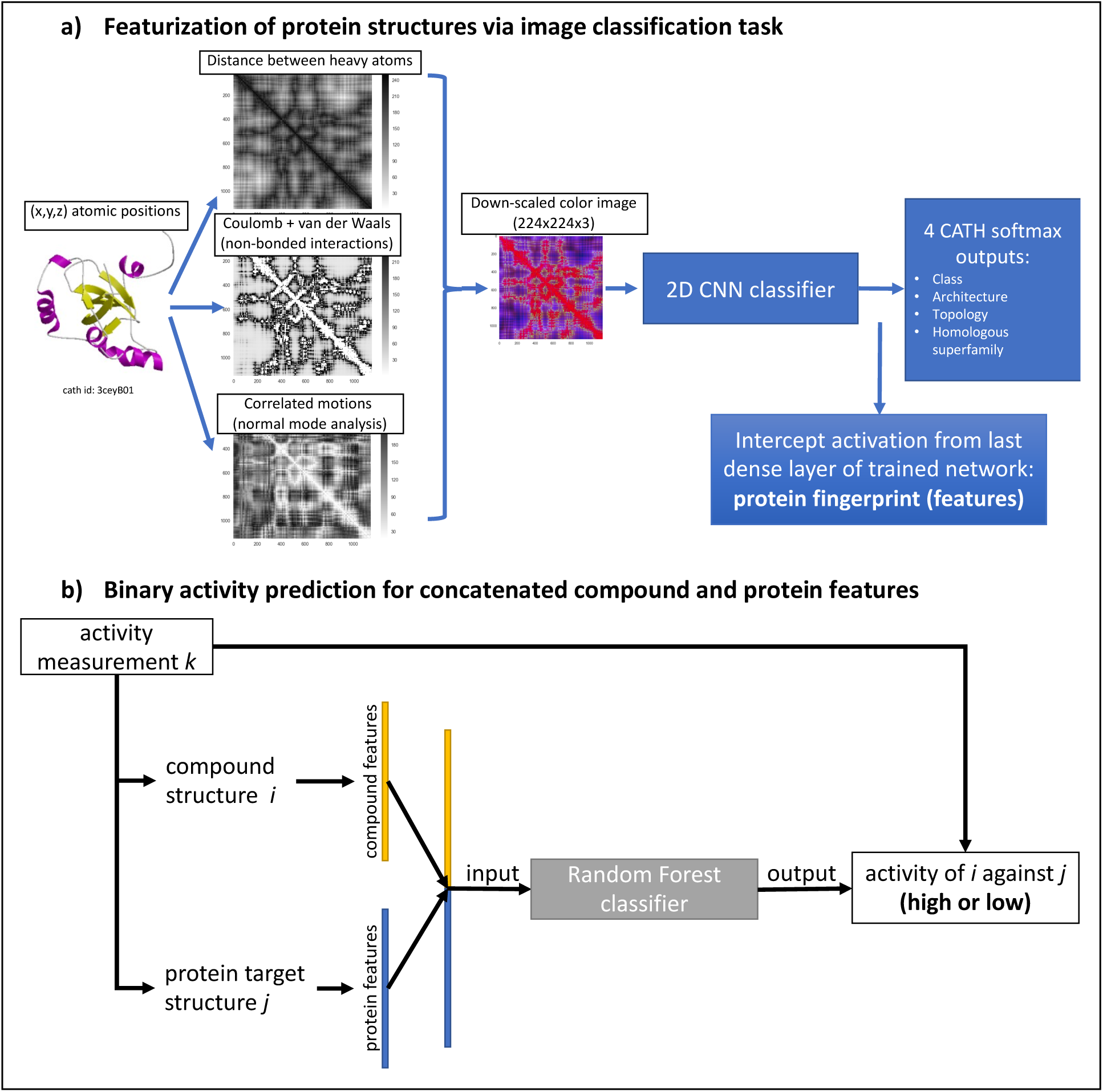
overview of machine learning workflow.

In its current version, the presented method, will scale down the 2D protein images from their individual native resolutions (the number of heavy atoms squared) to a common size such as 224×224 – which was used as the default size in many of the original publications of the deep learning models used here. The implications of this practice will be discussed in the Discussion section.

With slight modifications, successful image classification networks such as VGG16 can be used almost out-of-the-box, by simply changing the number of outputs to match the training data set at hand. In addition, the presented method works by connecting four parallel output layers to a newly added densely connected layer at the end of the network, one for each level of classification detail provided by CATH. This makes it effectively a multi-task learning network which has been shown to offer improvements in prediction [21]. Protein fingerprints (PFP) are simply the outputs of the last ReLU activation layer that are used as features by the connected softmax outputs to make a prediction. The length of a PFP is thus determined by the number of neurons in the last pre-output layer. Since changing this PFP length effectively also reduced network capacity, two different sizes of PFPs were tested to see any potential effect on training performance.

Table 1 summarizes training and validation performance of four different network architectures and two PFP sizes. Networks were trained until validation loss stopped improving and the number of epochs trained is given in the table as well. Training loss was at similarly low values for DenseNet121, Inception v3, and ResNet50, while VGG16 typically had one order of magnitude higher loss. The same is true for validation loss, where DenseNet121 based architectures showed the lowest values. Classification accuracies of the parallel tasks achieved always at least 99% during training and validation accuracy decreased with the number of classes per task with lowest values reaching 83-88% for “Homologous Superfamily” for three architectures, while VGG16 again underperformed. There is considerable over-fitting, the implications and potential remedies are discussed further below.

**Table 1:** Comparison of Deep Convolutional Neural Network architectures commonly used in supervised image classification problems applied to the encoding of protein feature vectors of length 128 and 512, trained on CATH. Cell coloring is applied per column and ranges from red (lowest value per column), over white, to green (highest value).

| architecture | PFP length | epochs trained | categorical crossentropy loss |  | "Class" accuracy 4 labels |  | "Architecture" accuracy 40 labels |  | "Topology" accuracy 1364 labels |  | "Homology" accuracy 2714 labels |  | PFP homogeneity score |  |  |  |
| --- | --- | --- | --- | --- | --- | --- | --- | --- | --- | --- | --- | --- | --- | --- | --- | --- |
|  |  |  | train | val | train C | val C | train A | val A | train T | val T | train H | val H | C | A | T | H |
| DenseNet 121 | 128 | 56 | 0.0300 | 2.1200 | 99.90% | 98.56% | 99.91% | 95.23% | 99.93% | 91.56% | 99.92% | 86.31% | 0.924 | 0.903 | 0.961 | 0.970 |
|  |  | 56 | 0.0252 | 2.1230 | 99.96% | 98.63% | 99.91% | 95.40% | 99.97% | 92.13% | 99.98% | 86.65% | 0.921 | 0.912 | 0.961 | 0.970 |
|  |  | 70 | 0.0221 | 2.1677 | 99.90% | 98.56% | 99.96% | 95.13% | 99.93% | 91.61% | 99.91% | 86.55% | 0.918 | 0.905 | 0.960 | 0.969 |
|  | 512 | 44 | 0.0053 | 2.0043 | 100.00% | 98.51% | 100.00% | 95.18% | 100.00% | 92.33% | 100.00% | 87.24% | 0.922 | 0.885 | 0.952 | 0.962 |
|  |  | 49 | 0.0039 | 2.0852 | 100.00% | 98.68% | 99.99% | 95.16% | 100.00% | 92.28% | 100.00% | 87.12% | 0.926 | 0.880 | 0.951 | 0.960 |
|  |  | 49 | 0.0051 | 2.0510 | 99.99% | 98.70% | 99.99% | 95.30% | 100.00% | 92.27% | 100.00% | 87.12% | 0.925 | 0.893 | 0.953 | 0.961 |
| Inception v3 | 128 | 49 | 0.0293 | 2.8683 | 99.90% | 97.90% | 99.97% | 93.63% | 99.94% | 88.56% | 99.89% | 84.74% | 0.212 | 0.298 | 0.652 | 0.740 |
|  |  | 40 | 0.0589 | 2.8501 | 99.90% | 97.96% | 99.87% | 93.59% | 99.86% | 88.25% | 99.82% | 84.24% | 0.269 | 0.263 | 0.615 | 0.718 |
|  |  | 60 | 0.0249 | 2.2118 | 99.91% | 98.64% | 99.96% | 95.48% | 99.92% | 91.92% | 99.95% | 86.47% | 0.926 | 0.897 | 0.963 | 0.973 |
|  | 512 | 38 | 0.0069 | 2.7395 | 99.99% | 97.82% | 100.00% | 93.55% | 100.00% | 88.73% | 99.98% | 85.01% | 0.042 | 0.243 | 0.653 | 0.746 |
|  |  | 47 | 0.0034 | 2.8140 | 100.00% | 97.68% | 100.00% | 93.51% | 100.00% | 88.70% | 100.00% | 84.87% | 0.105 | 0.201 | 0.619 | 0.727 |
|  |  | 42 | 0.0067 | 2.0951 | 99.99% | 98.59% | 100.00% | 95.52% | 100.00% | 92.07% | 99.98% | 86.69% | 0.924 | 0.885 | 0.957 | 0.967 |
| ResNet 50 | 128 | 53 | 0.0183 | 2.3034 | 99.98% | 98.35% | 99.96% | 93.99% | 99.96% | 90.47% | 99.98% | 84.53% | 0.917 | 0.896 | 0.956 | 0.966 |
|  |  | 49 | 0.0220 | 2.3435 | 99.92% | 98.34% | 99.97% | 94.29% | 99.94% | 90.13% | 99.98% | 83.88% | 0.920 | 0.902 | 0.958 | 0.965 |
|  |  | 71 | 0.0131 | 2.3876 | 99.94% | 98.44% | 100.00% | 94.12% | 99.99% | 90.17% | 99.95% | 84.29% | 0.917 | 0.894 | 0.957 | 0.967 |
|  | 512 | 40 | 0.0052 | 2.2145 | 100.00% | 98.19% | 99.98% | 94.05% | 100.00% | 90.64% | 100.00% | 84.88% | 0.915 | 0.866 | 0.948 | 0.958 |
|  |  | 43 | 0.0039 | 2.2289 | 100.00% | 98.15% | 100.00% | 93.99% | 100.00% | 90.72% | 100.00% | 85.26% | 0.913 | 0.879 | 0.949 | 0.959 |
|  |  | 62 | 0.0020 | 2.2210 | 100.00% | 98.26% | 100.00% | 94.11% | 100.00% | 90.96% | 100.00% | 85.16% | 0.918 | 0.885 | 0.949 | 0.958 |
| VGG16 | 128 | 53 | 0.0779 | 3.4529 | 99.82% | 97.31% | 99.74% | 90.59% | 99.74% | 84.60% | 99.57% | 76.69% | 0.894 | 0.811 | 0.937 | 0.948 |
|  |  | 54 | 0.1186 | 3.5379 | 99.75% | 96.74% | 99.69% | 90.34% | 99.58% | 83.94% | 99.25% | 76.21% | 0.887 | 0.794 | 0.932 | 0.942 |
|  |  | 54 | 0.1073 | 3.5075 | 99.66% | 96.96% | 99.54% | 90.31% | 99.74% | 84.94% | 99.33% | 76.63% | 0.885 | 0.797 | 0.932 | 0.943 |
|  | 512 | 48 | 0.4818 | 4.1244 | 99.19% | 96.03% | 95.44% | 85.08% | 97.37% | 79.50% | 96.42% | 69.23% | 0.587 | 0.620 | 0.888 | 0.906 |
|  |  | 48 | 0.0569 | 3.7016 | 99.98% | 96.26% | 99.91% | 89.30% | 99.86% | 83.82% | 99.81% | 75.95% | 0.867 | 0.704 | 0.906 | 0.923 |
|  |  | 56 | 0.1279 | 3.5749 | 99.88% | 96.96% | 99.21% | 89.66% | 99.66% | 84.23% | 99.32% | 75.55% | 0.742 | 0.654 | 0.906 | 0.922 |

As an alternative measure of performance, PFPs were clustered in their original dimension (128 or 512) using K-means where k was set to the four different numbers of classes per task in the outputs. The resulting cluster labels were then compared to the original CATH labels and the Homogeneity score as implemented in Scikit-Learn [22] was calculated where a 1.0 corresponds to a perfect overlap. This score allows to observe further performance differences across network architectures, but also between multiple instances of the same architecture (three models were trained for each architecture). Inception V3 showed considerable differences between replicates, showing some of the best and worst Homogeneity scores. DenseNet121 was the best-performing architecture across all metrics.

Larger PFPs of length 512 only provided slight improvements in training metrics potentially owed to the added network capacity but did not perform better in the homogeneity score than the architectures with PFPs of length 128.

Figure 2 provides a visualization of a 512-dimensional PFP space with PFPs predicted for all protein domains in the CATH data set. This distribution was reduced to two dimensions using t-SNE (t-distributed stochastic neighbor embedding) [23]. Colors indicate the CATH labels at first two classification levels (Class and Architecture). The model used to produce these fingerprints are indicated by an asterisk (*) in Table 1 and corresponds to the model with lowest overall validation loss. Overall, visual inspection of Figure 2 reveals a clustering of same-color data points which is consistent with the high Homogeneity scores. Some visual imperfections can be identified in the left panel of Figure 2 where four large clusters are expected to be separated, but visually some discontinuities can be seen. Given the high Homogeneity scores across CATH levels, this imperfection could be due to the stochasticity of t-SNE. Another DenseNet121 model (with PFP length 128) had very slightly higher score at the Class-level and also has a more homogeneous appearance in the t-SNE plot (not shown).

**Figure 2:**
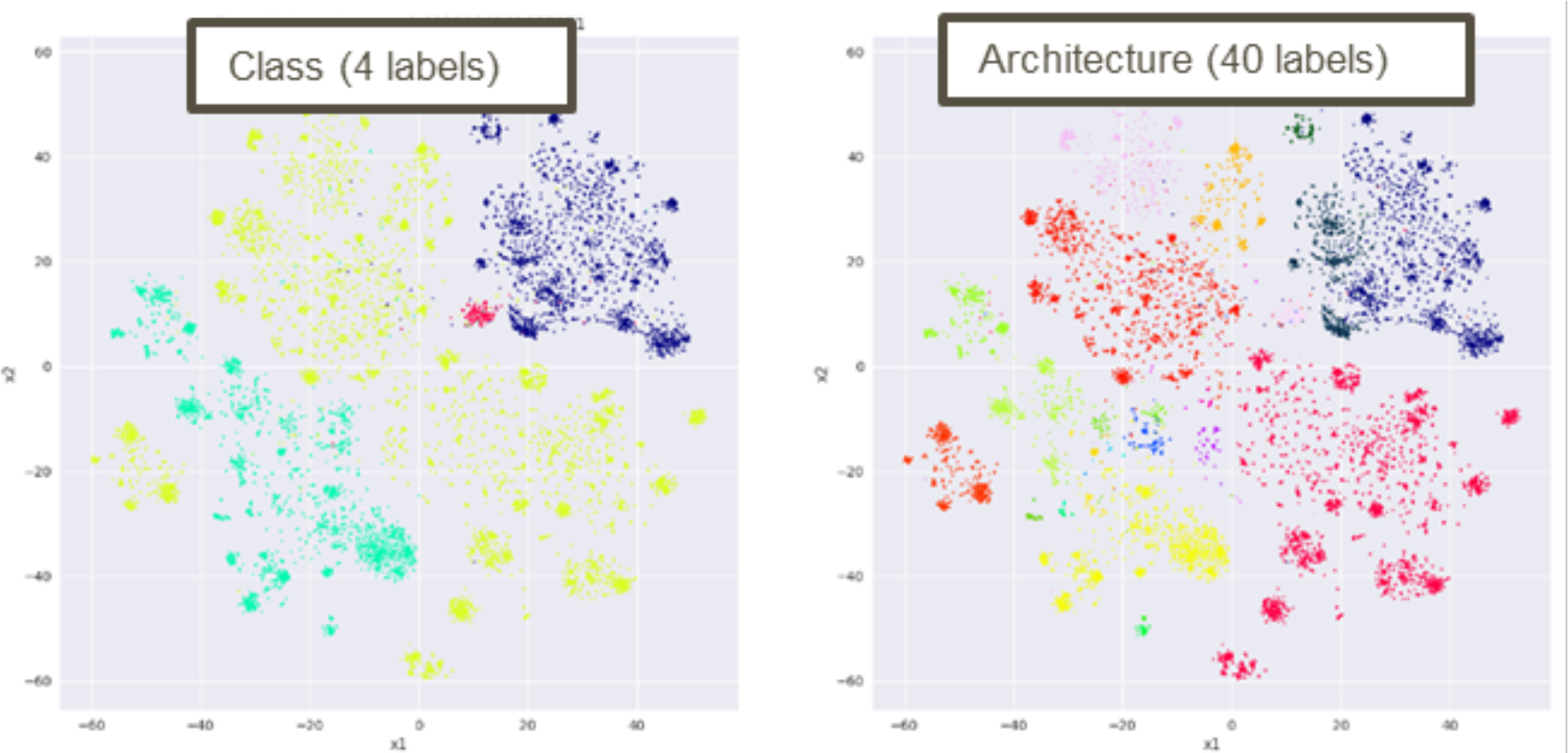
Dimensionality reduction of 512-dimensional protein feature vectors obtained from a DenseNet architecture to 2D via t-SNE. All protein domains in the CATH non-redundant data set are colored based on ground truth labels of the Class and Architecture outputs.

In summary, a 128-dimensional fingerprint can be sufficient to achieve a near perfect encoding of the CATH data set. This is a stark contrast to the amount of information used as input, i.e. the 224×224×3 = 150528 color intensity values of the 2D color images. This number of features is an impediment to any useful analysis and t-SNE plots on the raw image values convey the inability to identify meaningful clusters in the same data set of CATH domains (not shown).

While Table 1 only lists supervised deep learning approaches, there are also successful other approaches used on images, such as the U-net architecture for image segmentation, i.e. the task of outlining objects belonging to a known category within an image. The U-net architecture essentially follows the structure of a deep, multi-layer auto-encoder with skip connections from the encoder to the decoder arm of the network [20]. The U-net architecture was used here with the modification of reconstructing the input image instead of a binary / multi-label segmentation mask. This version of the U-net is therefore truly unsupervised in the sense that no additional training labels were needed. Figure 3 shows the same t-SNE map as Figure 2, but based on fingerprints extracted from the activations of the 1024-dimensional bottleneck layer of U-net (after mapping each of the 1024 2D feature maps to single averages). In the absence of CATH training labels, the fingerprints do not cluster visually when coloring data points according to CATH labels. Nevertheless, the U-net fingerprints performed at the same level as other fingerprints in the activity prediction task (see below).

**Figure 3:**
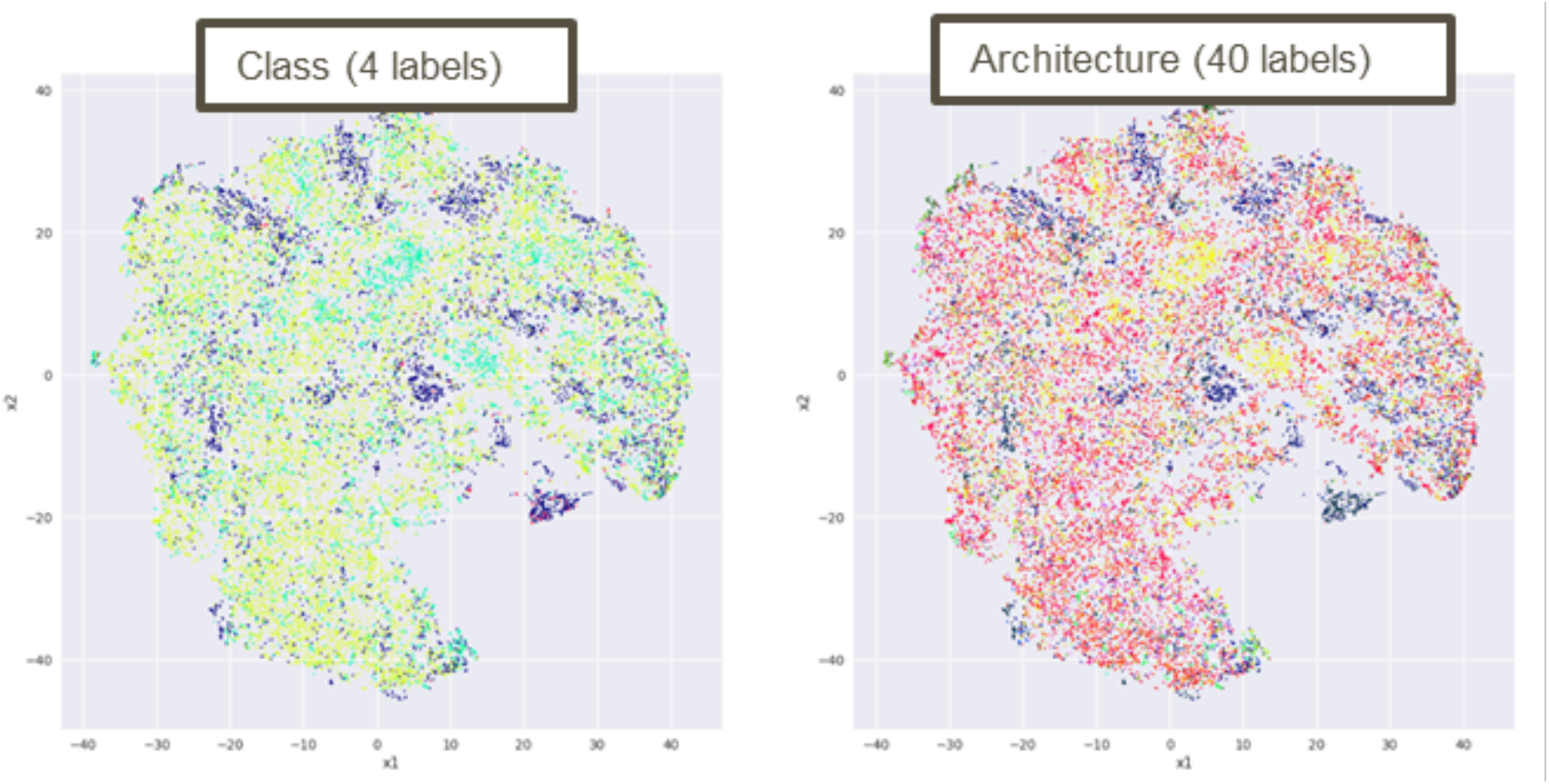
Dimensionality reduction of 1024-dimensional protein feature vectors obtained from a U-Net architecture to 2D via t-SNE. All protein domains in the CATH non-redundant data set are colored based on ground truth labels of the Class and Architecture outputs.

### Protein fingerprints applied to ChEMBL activity prediction

After encoding the CATH dataset into 128 and 512-dimensional fingerprints (1024 in the case of U-net), the general applicability of such fingerprints was demonstrated on the ChEMBL (version 23) dataset to predict chemical activity of small molecule compounds against specific proteins measured experimentally and recorded as the half-maximal concentration of the compound at which the anticipated effect was observed, e.g. inhibition of protein activity. ChEMBL provides a standardized value for various types of activity and calls it the p-ChEMBL value. The Methods section describes how the ChEMBL database was used to generate an approximately balanced binary classification dataset without trivial class assignment (e.g. one target always inactive) and how the single-domain target protein was assigned to the PDB code of the best-matching structure which in turn was then converted to a distance-based representation (“image”) and encoded to a fixed-length fingerprint using the CATH-trained classifier described above.

Each activity entry in this ChEMBL dataset thus provides structural information of both the compound and the protein, but independently of each other, i.e. without knowing the exact binding position or mechanism. Compounds were encoded using two alternative fingerprint representations commonly used and implemented in the chemo-informatics software RDkit [24], MACCS (167 bits) and Morgan (1024 bits, radius=3), see [25] for a comparison of different fingerprints. With this compound fingerprint (CFP) and the corresponding protein fingerprint (PFP) concatenated, the final feature vector was used as input to another machine learning model. Here, a random forest ensemble classifier was chosen due to its good performance, generalization metrics, and interpretability. However, any other classifier may be used in its stead. A binary classification setup was chosen to predict whether a CFP and PFP combination falls into the ‘high’ or ‘low’ activity bin – as this task is more attainable than an exact regression predictor of actual activity (p-ChEMBL) values, especially since these measurements come from very different types of assays. Since activity datasets like ChEMBL can be biased towards certain compound structures or targets – providing a lot of data for some, but little for others – two alternative feature vectors were created for comparison. First, the CFP alone was used as input for activity prediction. In this case, only the compound structure itself can provide any information that would lead to a higher-than-chance accuracy of predicting activity and would indicate that certain compound structural features are over-represented in the datasets. Second, the CFP was concatenated with a one-hot encoded array indicating which of the targets the compound was measured against. In this case, predictions will be improved if certain compounds or compound classes have been tested on particular targets more than others and if certain targets can be associated with either high or low activity. Third, CFP and PFP are concatenated, providing structure-level details of both. Improvements to the prediction scores would mean that the structural information of the protein contains additional clues on activity when combined with the CFP. PFPs were generated using the DenseNet121 architectures with lowest validation loss and with PFP lengths 128 and 512.

Table 2 shows the prediction results of the ChEMBL activity classifier. Looking at the mean performance in a 5-fold stratified cross-validation (CV) setup, all metrics indicate only a slight improvement over a random baseline of 0.5 when just using the CFP with no protein features. Adding the one-hot encoded index of the target on the other hand, provides an enormous performance boost. Such an approach can serve as a baseline model that mostly utilizes hidden biases in the way certain types of molecules are associated with certain types of targets and/or activity patterns – rather than making use of molecular data of each individual target protein. Adding structure-based PFPs provides improved prediction metrics over the CFP+index approach and at a much lower number of features. PFPs also perform at the same level as sequence-derived word embeddings via ProtVec.

**Table 2:** Training metrics of ChEMBL activity prediction using a binary random forest classifier. Each compound and protein feature label contains the number of features at the end. Bold protein feature labels indicate contributions from this paper. Tables are sorted by decreasing OOB score.

| compound features | compound feature importance | protein feature importance | protein features | oob score | best score f1 |
| --- | --- | --- | --- | --- | --- |
| Morgan3-1024 | 0.6213 | 0.3787 | DenseNet-128 | 0.8457 | 0.8280 |
| Morgan3-1024 | 0.6133 | 0.3867 | DenseNet-512 | 0.8449 | 0.8285 |
| Morgan3-1024 | 0.6176 | 0.3824 | U-Net-1024 | 0.8447 | 0.8272 |
| Morgan3-1024 | 0.6183 | 0.3817 | ProtVec-300 | 0.8446 | 0.8263 |
| MACCS-166 | 0.6078 | 0.3922 | DenseNet-128 | 0.8406 | 0.8234 |
| MACCS-166 | 0.5962 | 0.4038 | ProtVec-300 | 0.8398 | 0.8223 |
| MACCS-166 | 0.5867 | 0.4133 | DenseNet-512 | 0.8397 | 0.8229 |
| MACCS-166 | 0.6202 | 0.3798 | U-Net-1024 | 0.8391 | 0.8220 |
| MACCS-166 | 0.5506 | 0.4494 | onehot-target-871 | 0.8149 | 0.7980 |
| Morgan3-1024 | 0.5861 | 0.4139 | onehot-target-871 | 0.8117 | 0.7914 |
| MACCS-166 | 1.0000 | 0.0000 | None | 0.6690 | 0.5973 |
| Morgan3-1024 | 1.0000 | 0.0000 | None | 0.5987 | 0.5673 |

b) Compound group 5-fold cross-validation (groups are 5 k-means clusters of compound features)
| compound features | compound feature importance | protein feature importance | protein features | oob score | best score f1 |
| --- | --- | --- | --- | --- | --- |
| Morgan3-1024 | 0.6214 | 0.3786 | DenseNet-128 | 0.8458 | 0.7612 |
| Morgan3-1024 | 0.6131 | 0.3869 | DenseNet-512 | 0.8454 | 0.7604 |
| Morgan3-1024 | 0.6174 | 0.3826 | U-Net-1024 | 0.8448 | 0.7613 |
| Morgan3-1024 | 0.6184 | 0.3816 | ProtVec-300 | 0.8442 | 0.7619 |
| MACCS-166 | 0.6111 | 0.3889 | DenseNet-512 | 0.8402 | 0.7150 |
| MACCS-166 | 0.6081 | 0.3919 | DenseNet-128 | 0.8401 | 0.7173 |
| MACCS-166 | 0.6121 | 0.3879 | ProtVec-300 | 0.8396 | 0.7153 |
| MACCS-166 | 0.6203 | 0.3797 | U-Net-1024 | 0.8390 | 0.7166 |
| MACCS-166 | 0.5509 | 0.4491 | onehot-target-871 | 0.8149 | 0.7001 |
| Morgan3-1024 | 0.5848 | 0.4152 | onehot-target-871 | 0.8120 | 0.7495 |
| MACCS-166 | 1.0000 | 0.0000 | None | 0.5916 | 0.4583 |
| Morgan3-1024 | 1.0000 | 0.0000 | None | 0.5898 | 0.5178 |

c) Protein group 5-fold cross-validation (groups are 5 k-means clusters of protein features)
| compound features | compound feature importance | protein feature importance | protein features | oob score | best score f1 |
| --- | --- | --- | --- | --- | --- |
| MACCS-166 | 0.1110 | 0.8890 | DenseNet-128 | 0.7698 | 0.6291 |
| Morgan3-1024 | 0.1038 | 0.8962 | ProtVec-300 | 0.7684 | 0.6338 |
| MACCS-166 | 0.0936 | 0.9064 | DenseNet-512 | 0.7683 | 0.6142 |
| MACCS-166 | 0.0949 | 0.9051 | ProtVec-300 | 0.7676 | 0.6280 |
| Morgan3-1024 | 0.1055 | 0.8945 | DenseNet-512 | 0.7672 | 0.6410 |
| Morgan3-1024 | 0.1130 | 0.8870 | DenseNet-128 | 0.7656 | 0.6385 |
| MACCS-166 | 0.0777 | 0.9223 | U-Net-1024 | 0.7646 | 0.6525 |
| Morgan3-1024 | 0.0942 | 0.9058 | U-Net-1024 | 0.7637 | 0.6362 |
| MACCS-166 | 1.0000 | 0.0000 | None | 0.6698 | 0.5241 |
| MACCS-166 | 0.3289 | 0.6711 | onehot-target-871 | 0.6649 | 0.6059 |
| Morgan3-1024 | 0.2950 | 0.7050 | onehot-target-871 | 0.6596 | 0.6119 |
| Morgan3-1024 | 1.0000 | 0.0000 | None | 0.5900 | 0.3576 |

Group-based 5-fold CV requires the model to learn from 4 out of 5 clusters of features while predicting the 5^th^ cluster. Two different group splits were explored here, one where clusters were formed based on the compound features, and one based on the PFPs. Feature importances were summed up for all compound features and all protein features, separately. It was found that compound-based group splits resulted in lower F1 accuracy than stratified splits, but compound features contributed around 60% to overall feature importance in both types of splits. Splitting according to PFP groups, on the other hand, dropped compound feature importance to around 10%, i.e. 90% of feature importance came from the proteins. At the same time, the features making use of molecular-level information (either sequence or structure) performed at a much higher level than the one-hot encoded target index, whereas one-hot encoding was sufficient to achieve relatively high cross-validation scores in the other two CV strategies. A proper test set of data not involved in either training or validation, however, was not used here.

Scikit-Learn’s GridSearchCV function was used for CV-based hyper-parameter search, finding better performing settings for some of the parameters specific to random forest models, namely the number of estimators in the ensemble, the maximum number of input features and the maximum tree depth. Results are only shown from models that achieved the highest F1 score for a given set of input features. The F1 score is a balanced accuracy that combines both precision and recall. The OOB (out of bag) score provided an additional metric of generalization for random forest models, where randomly excluded samples from the training set are used to validate predictors.

## DISCUSSION

An efficient reduction of protein structures to a small, consistent set of features (fingerprint) is of high importance in Structural Bioinformatics and allows for an accurate estimation of protein relationships and may be useful in activity prediction of small molecules.

Here, a new machine learning approach showed that existing image classification neural networks can be adopted to encode a protein fingerprint and that such a fingerprint seems to provide molecular detail that is comparable to the protein sequence level.

Models trained with CV on held-out protein feature clusters seem to make better use of the molecular detail of the proteins in the training set and use those features for enhanced generalization to unseen targets in the validation set. This is encouraging, especially in the light of reported cases where biases in the compound structures were sufficient for high model performance and detailed 3D structure information was not utilized by the model at all [26]. However, it seems that at the level of image resolution used here, sequence-derived features (ProtVec) performed just as well.

The approach outlined here still has several limitations to overcome. Most importantly, the practical requirement of current deep learning frameworks to provide fixed-dimensional data batches discourages the use of full atomic resolution of proteins with very different sizes. Also, the limitation of GPU RAM currently necessitates to work with scaled-down images. However, time is likely to alleviate such limitations in the future in combination with more sophisticated neural network architectures and more powerful computer hardware. There is also no established method for data augmentation of proteins – a technique that is frequently used on images where small perturbations such as rotation, translation, and rescaling are used to artificially provide more training data and that allows the model to learn invariances over such perturbations.

The degree of over-fitting observed in Table 1 still suggests that this work would benefit from more data in the future, such as the full CATH domain set. The capacity of the neural networks used may also have been higher than required for the classification task, and indeed smaller PFPs (i.e. the last dense layer) performed as well or better than larger ones.

CATH protein classification may also not be the most challenging task that makes use of atomic detail. Future models should directly train neural networks on the input images while predicting molecular binding. A more thorough comparison between 1D, 2D, 3D and graph representations of proteins should be compared on the same benchmark. Most recently, graph convolutional networks have been applied on both small molecules and proteins and may be a more efficient way of representing both small and large molecules [27]. Such future benchmarks should also investigate the merits of including the non-bonded and dynamics-based inputs channels explored here, and for various tasks.

Activity prediction remains a challenging task, not least because of the likely issues present in the commonly used data sets such as ChEMBL, which for the most part are collections of experiments conducted independently of each other and inconsistently. The mapping from assay to protein sequence/structure is also not always possible with certainty so that it is very likely that false labels were present in the data set used here. 3D structures are also not available nearly as abundantly as protein sequences.

Another limitation of all current machine learning models is that they can either use the protein sequence or the 3D structure, but not both. Ideally, models that featurize proteins should work with just the sequence as input but produce features consistent with 3D information and thus need to be trained on both modalities in some way.

## ACKNOWLEDGEMENTS

Thanks to numerous members of the computational sciences and AI/ML teams at GSK for valuable discussions.

## Notes

#### Summary of Updates

Parts of Figure 1 were missing due to the PDF conversion and have now been restored.

